# Methodological considerations when assessing mitochondrial respiration and biomarkers for mitochondrial content in human skeletal muscle

**DOI:** 10.1101/2021.03.25.436899

**Authors:** Jujiao Kuang, Nicholas J Saner, Javier Botella, Matthew J-C Lee, Cesare Granata, Zhenhuan Wang, Xu Yan, Jia Li, Amanda J Genders, David J Bishop

## Abstract

**Background:** The assessment of mitochondrial respiration and mitochondrial content are two common measurements in the fields of skeletal muscle research and exercise science. However, to verify the validity of the observed changes in both mitochondrial respiration and mitochondrial content following an intervention such as exercise training, it is important to determine the reliability and reproducibility of the experimental design and/or techniques employed. We examined the repeatability of widely used methodologies for assessing mitochondrial respiration and mitochondrial content, respectively; the measurement of maximal mitochondrial oxidative phosphorylation in permeabilized muscle fibres using high-resolution respirometry, and the measurement of citrate synthase activity as a biomarker for mitochondrial content in a microplate with spectrophotometer.

**Result:** For mitochondrial respiration, the coefficient of variation for repeated measurements using muscle sampled from same biopsy decreased from 12.7% to 11% when measured in triplicate with outliers excluded, rather than in duplicate. The coefficient of variation was 9.7% for repeated muscle biopsies sampled across two separated days. For measurements of citrate synthase activity, the coefficient of variation was 3.5% of three technical repeats on the same plate, 10.2% for duplicate analyses using the same muscle lysate when conducted in the same day, and 30.5% when conducted four weeks apart.

**Conclusion:** We have provided evidence for important technical considerations when measuring mitochondrial respiration with human skeletal muscle: 1) the relatively large technical variability can be reduced by increasing technical repeats and excluding outliers; 2) the biological variability and absolute mitochondrial respiration value of the participants should be considered when estimating the required sample size; 3) a new threshold of 15% for the increase in respiration rate after the addition of cytochrome *c* test for testing mitochondrial outer membrane integrity. When analysing citrate synthase activity, our evidence suggests it is important to consider the following: 1) all samples from the same study should be homogenized and measured at the same time using the same batch of freshly made chemical reagents; 2) biological variability should be considered when detecting small change in mitochondrial content; 3) the relative change should be used to compare the outcomes from different studies.

## BACKGROUND

Found in most eukaryotic cells, mitochondria are membrane-enclosed organelles that generate adenosine triphosphate (ATP), the primary energy molecule of metabolism, in a process termed oxidative phosphorylation (OXPHOS). Mitochondria are highly sensitive and adaptive to a range of environmental stresses [1], including exercise training [2]. A key adaptation to exercise is the process of mitochondrial biogenesis, which is defined as “the making of new components of the mitochondrial reticulum” [3]. This process can be reflected by changes in mitochondrial respiration (measured via the rate of oxygen consumption) and by changes in mitochondrial content (either measured directly using transmission electron microscopy (TEM) or via markers such as citrate synthase (CS) activity [4]). There are many studies that demonstrate increases in these measures following exercise training (reviewed in [5]); thus, both have become useful and commonly used measurements of training-induced mitochondrial adaptions in human skeletal muscle.

It is often incorrectly assumed that exercise training simultaneously promotes similar changes in mitochondrial respiration and content. However, many studies show that exercise training can increase mitochondrial respiration or mitochondrial content independently [6–10]. Thus, it has been recommended to measure both when examining mitochondrial adaptations to exercise training [11]. However, to ensure the accurate assessment, it is imperative to understand and determine the reliability of these commonly measured mitochondrial adaptations.

The classic method of assessing mitochondrial respiration involves measuring the oxygen consumption rate using an oxygen-sensitive electrode [12, 13]. High-resolution respirometers, such as the Oxygraph-2k respirometer (Oroboros, Innsbrunk, Austria), are commercially available and commonly used systems to assess mitochondrial respiration in yeast, cells, and tissue, including human skeletal muscle. Both isolated mitochondria and permeabilized muscle fibres can be used with this system [14]. However, although the use of isolated mitochondria provides less variation between technical repeats, as it results in a homogenous lysate, it also causes damage to, and loss of, mitochondria [15]. In contrast, using permeabilized muscle fibres allows the structural integrity of myocytes and the outer mitochondrial membrane to be better maintained, and preserves functional clusters of mitochondria and other cell organelles [16]. In addition, the use of permeabilized muscle fibres requires a smaller muscle sample. Consequently, the use of permeabilized muscle fibres with high-resolution respirometry has become the most common method to measure mitochondrial respiration in human skeletal muscle [6, 17, 18]. However, despite its widespread use in human exercise training studies ([5, 19–31], and several studies have been conducted to assess factors that might cause technical and/or biological variation [32–34], uncertainty remains regarding the reliability of this method. As such, additional research is needed to establish the reliability of mitochondrial respiration measures using permeabilized muscle fibres with high-resolution respirometry.

The measurement of mitochondrial volume density using TEM is considered the “gold standard” for determining mitochondrial content [35]. However, this analysis is expensive, time consuming, and it is often not readily available. Instead, the use of CS activity as a biomarker for mitochondrial content has become a standard approach in human exercise training studies [4] given its strong correlation with mitochondrial volume density (as measured by TEM) [4, 36], and being a less time-consuming technique. However, it is difficult to directly compare CS activity assay results between studies due to different protocols utilized and variations in how the results have been reported [37]. For instance, this analysis can be performed using freshly homogenized skeletal muscle [6] or freeze-dried muscle dissected free of visible blood and connective tissue [7]. Furthermore, the reaction can be carried out in a cuvette [38] or using a microplate with a plate reader [39].

Despite the widespread use of both measurements within the field of human exercise studies, some important questions remain unanswered. When assessing mitochondrial respiration (in permeabilized muscle fibres), the following factors remain undetermined: 1) the number of technical repeats that should be used; 2) whether the variability is affected by the absolute mitochondrial respiration value and muscle mass; what should be considered a valid cut-off for the mitochondrial membrane integrity test; and 4) what is the variability between repeated biopsies over a control period. When assessing mitochondrial content in human skeletal muscle (the CS activity assay), it is unclear: 1) if one can directly compare results derived using the same protocol on two separate occasions; 2) what is the variability in mitochondrial content between repeated biopsies over a control period; 3) whether it is appropriate to compare results derived from two different protocols (e.g., using a cuvette or microplate). Therefore, to obtain the most reproducible and accurate findings, the aim of the present study was to carefully examine the influence of such methodological variables and to provide guidance to the large number of researchers using these techniques.

## RESULTS

### Experiment 1: Reliability of mitochondrial respiration between the two chambers of the Oxygraph O2k in a yeast cell suspension

We assessed the variability of repeated measurements from individual chambers on the same respirometer, using freshly prepared yeast cell suspensions. This was done to minimize the between-sample biological variability occurring in human skeletal muscle samples. The respiration rates were consistent between the two chambers (*n* = 8; *P* = 0.484; Figure 1A and Supplementary Table 1) and were correlated (R = 0.798, *P* = 0.018; Figure 1B). The average coefficient of variation (CV) for the pairs of measurements between two chambers was 6.1% (range from 2.2% to 9.4%; Figure 1C), and the technical error of measurement (TE_M_) was 0.1 nmol·s^-1^·mg dry yeast^-1^ or 6.3% (Supplementary Table 1).

**Figure 1:**
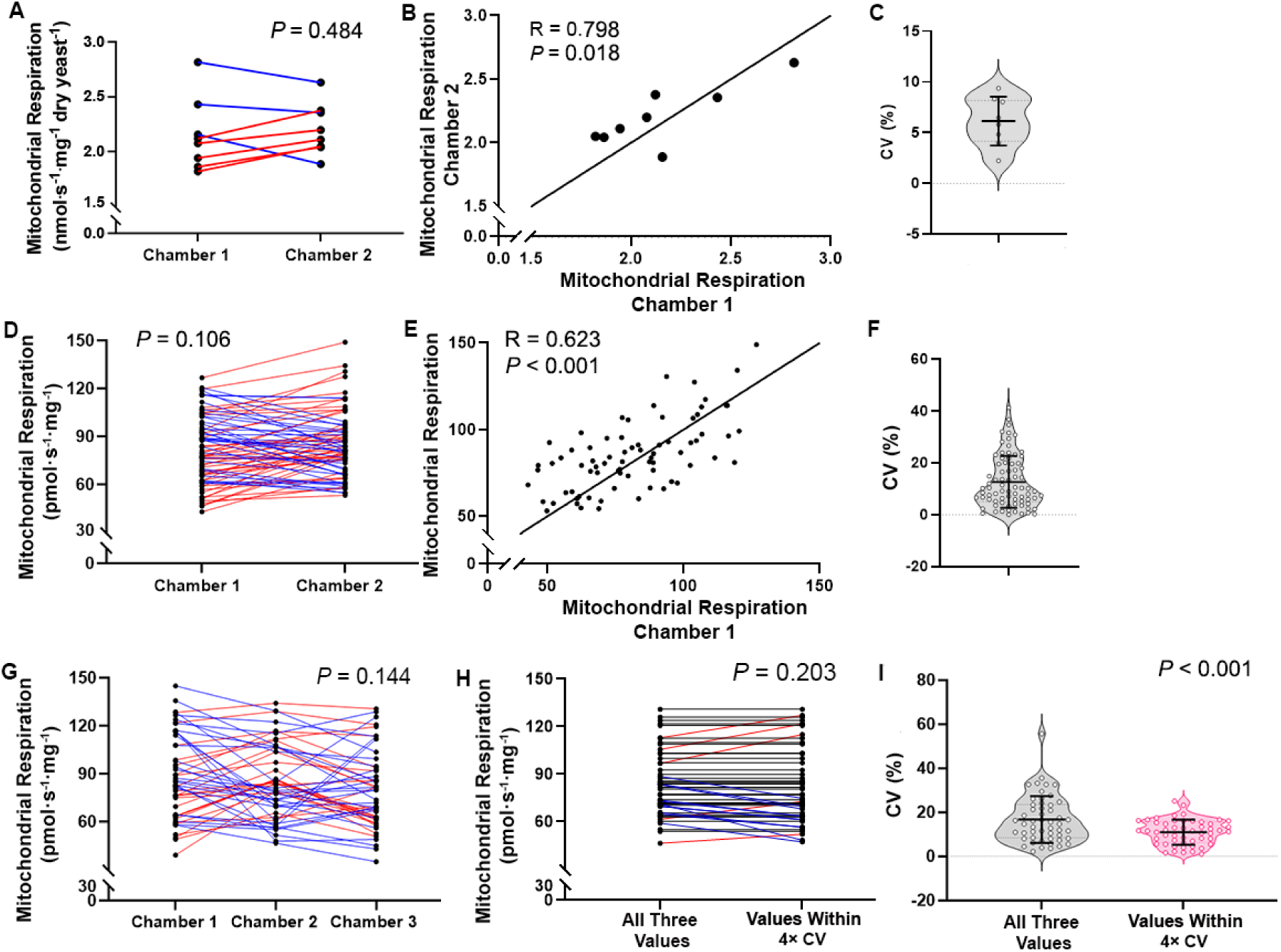
Reliability of mitochondrial respiration between chambers of Oxygraph O2k. **A, D, and G:** Mitochondrial respiration of yeast cell suspension (A, *n* = 8) using glucose as substrate, maximal mitochondrial respiration (CI + CII_p_) of permeabilized skeletal muscle (D, *n* = 76) measured in two chambers of the same respirometer, and permeabilized skeletal muscle measured in three chambers across two respirometers (G, *n* = 51). Individual measurements of the same sample are represented by black dots and connected with a line. The red lines represent an increase, whereas blue lines represent a decrease related to Chamber 1. **B, and E**: Pearson’s correlation between the two measurements of (B) yeast cell suspension and (E) permeabilized skeletal muscle. Values are in nmol·s-^1^·mg dry yeast^-1^ or (E) pmol·s-1·mg-1. Solid line shows the line of identity. **C, F, and I.** The respective coefficients of variation between two chambers of the same respirometer in (C) yeast cell suspension,(F) permeabilized skeletal muscle, and (I) the “All Three Values” (all three chambers), and “Values Within 4× CV” (excluding outliers outside of 4x the coefficients of variation) measurements of permeabilized skeletal muscle measured in three chambers across two respirometers. The coefficients of variation are represented by open circles; the mean and standard deviation are presented by the black line and error bars, respectively. The width of the shaded area of violin plot represents the proportion of the data located there. **H.** The two ways to calculate maximal mitochondrial respiration (CI + CII_p_) of permeabilized skeletal muscle measured in three chambers (data from G): all values from three chambers (“All Three Values”); or after excluding outlier outside of 4x the coefficients of variation (“Values Within 4× CV”). Individual respiration rates calculated for the same sample are represented by black dots and connected with a line. The black lines represent no change, red lines represent an increase, whereas blue lines represent a decrease related to “All Three Values”.

### Experiment 2: Reliability of mitochondrial respiration between the two chambers of the same Oxygraph O2k in permeabilized skeletal muscle fibre

Maximal mitochondrial respiration rates (CI + CII_p_) of the same permeabilized skeletal muscle samples (*n* = 76) measured in two separate chambers on the same respirometer were not significantly different from each other (*P* = 0.106; Figure 1D and Supplementary Table 1), and the two values were correlated (R = 0.623, *P* < 0.001; Figure 1E). The average CV of measurements between two chambers was 12.7% (range from 0.03% to 41.3%; Figure 1F), and the TE_M_ was 12.8 pmol·s^-1^·mg^-1^ or 15.4% (Supplementary Table 1).

### Experiment 3: Reliability of mitochondrial respiration between three chambers, across two Oxygraph O2k, in permeabilized skeletal muscle fibre

Data from 51 skeletal muscle biopsy samples were each analysed in triplicate simultaneously using three chambers across two respirometers (Supplementary Table 1, Figure 1G). The maximal mitochondrial respiration (CI + CII_p_) was calculated using the mean of the values from all three chambers (“All Three Values”; *mean ± SD*: 84.2 ± 21.0 pmol·s^-1^·mg^-1^; Figure 1H), or the mean of the values after excluding outliers (maximum of one outlier per sample) (“Values Within 4× CV”; *mean ± SD*: 82.9 ± 23.8 pmol·s^-1^·mg^-1^; Figure 1H). As reference, four times the CV of yeast respiration (representing the background variability of the instrument and operator combination) from the mean of three values was chosen as an acceptable range for normal data distribution, and any values outside of this range were excluded from data analysis due to their abnormal distance from the other values. The maximal respiration values were not significantly different between the three chambers (*P* = 0.144; Figure 1G), or between the two calculation methods (the “All Three Values” and “Values Within 4× CV”, *P* = 0.203; Figure 1H). The CV of measurements from the “All Three Values” and “Values Within 4× CV” were significantly different from each other (*P* < 0.001, Figure 1I): “All Three Values” = 16.7% (range from 2.2% to 55.6%); “Values Within 4× CV” = 11.0% (range from 1.0% to 25.1%). The TE_M_ between the “All Three Values” was 15.2 pmol·s^-1^·mg^-1^ or 18.1%. The TE_M_ between “Values Within 4× CV” was 9.8 pmol·s^-1^·mg^-1^ or 11.7%.

### The effect of absolute mitochondrial respiration value on reliability of mitochondrial respiration between chambers

Based on the results from Experiments 2 and 3, the relationship between the absolute mitochondrial respiration value and the variation between duplicate measurements was investigated. In human exercise training studies where mitochondrial respiration rate was analyzed in permeabilized skeletal muscle fibres with high-resolution respirometry, most of the reported maximal respiration rates of healthy moderately trained male participants before training was 75 to 100 pmol·s^-1^·mg^-1^, and some of the studies reported higher respiration rates after training [6, 9, 36, 40–42]. Therefore, our maximal mitochondrial respiration values (CI + CII_p_) were split into three groups: ‘*Low’* (< 75 pmol·s^-1^·mg^-1^), ‘*Medium’* (75 to 100 pmol·s^-1^·mg^-1^), and ‘*High’* (> 100 pmol·s^-1^·mg^-1^). The CV of the data from Experiment 2 (*n* = 76, Figure 2A) and Experiment 3 (*n* = 51, 3 chambers with no outlier excluded, Figure 2B) were calculated in each group. There was no significant difference between the CV of the three groups in Experiment 2 (*P* = 0.096; Figure 2A); however, in Experiment 3, the CV were significantly different between the three groups (*P* = 0.030; Figure 2B). In Experiment 2, the absolute TE_M_ for the three groups were similar: Low (13.1 pmol·s^-1^·mg^-1^); Medium (12.8 pmol·s^-1^·mg^-1^); and High (12.4 pmol·s^-1^·mg^-1^) value groups. However, as the mean respiration values were different between the three groups, the relative TE_M_ decreased from 20.3% (Low) to 14.9% (Medium) and 11.0% (High). In Experiment 3, the relative TE_M_ also decreased with absolute maximal mitochondrial respiration rates, from 21.7% (Low, TE_M_: 14.2 pmol·s^-1^·mg^-1^) to 17.0% (Medium, TE_M_: 14.8 pmol·s^-1^·mg^-1^) and 13.0% (High, TE_M_: 15.0 pmol·s^-1^·mg^-1^).

**Figure 2:**
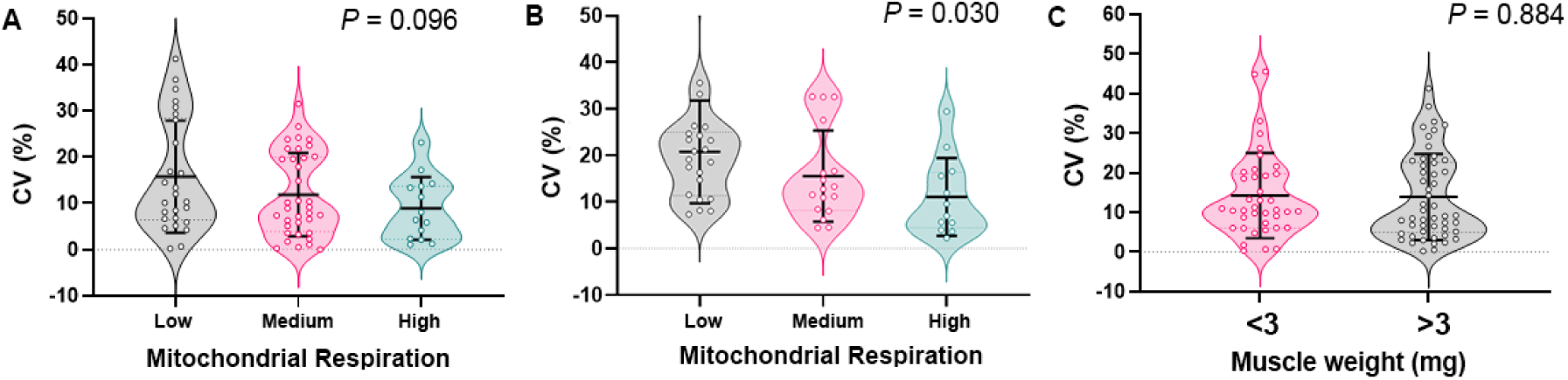
Reliability of mitochondrial respiration between two measurements in relation to absolute respiration rate and muscle mass. **A and B**: Coefficient of variation between maximal mitochondrial respiration rates (CI + CII_p_) of permeabilized skeletal muscle measured in (A) two chambers (*n* = 76, from Experiment 2) and (B) the three chambers from the same permeabilized skeletal muscle samples (*n* = 51, no data excluded, from Experiment 3), categorized according to low (< 75 pmol·s^-1^·mg^-1^), medium (75 to 100 pmol·s^-1^·mg^-1^), and high respiration rates (> 100 pmol·s^-1^·mg^-1^). **C**: Coefficient of variation of maximal mitochondrial respiration (CI + CII_p_) in permeabilized skeletal muscle (*n* = 89 pairs, samples from Experiments 2 and 3), according to sample mass (< or > 3 mg). Individual coefficient of variation are represented by open circles; the mean and standard deviation are presented by the black line and error bars, respectively. The width of the shaded area of violin plot represents the proportion of the data located there.

### The effect of muscle mass on reliability of mitochondrial respiration between chambers

From Experiments 2 and 3, 89 pairs of measurements, for which both chambers had similar muscle mass, were selected and categorised into two groups: (1) both chambers with muscle mass < 3 mg or (2) > 3 mg. Regardless of the muscle sample mass, there was no significant difference in the CV between groups (*P* = 0.884; Figure 2C), nor the TE_M_ (< 3 mg: 14.6 pmol·s^-1^·mg^-1^ or 17.5%; > 3 mg: 14.5 pmol·s^-1^·mg^-1^ or 17.4%).

### Validation of cytochrome *c* test

The addition of cytochrome *c* to the respiration chamber should be used to assess mitochondrial outer membrane integrity, and a 10% or more rise in respiration after the addition of cytochrome *c* is often considered as indicative of loss of outer membrane integrity [43]. The validity of this threshold was assessed using 31 muscle samples analyzed in triplicate (*i.e.*, three chambers), where one chamber elicited a > 10% increase in mitochondrial respiration rate after adding cytochrome *c* (the often-used criteria for “Failed”), and the other two elicited a < 10% increase in mitochondrial respiration after the addition of cytochrome *c* (the often-used criteria for “Passed”; Figure 3A). The samples were grouped according to the increase in respiration of their respective “Failed” chamber (10 to 15%, 15 to 20%, or > 20%); the maximal mitochondrial respiration rates (CI + CII_p_) of the pair of “Passed” and corresponding “Failed” chambers, recorded before the addition of cytochrome *c*, were then compared. There was a significant difference in respiration values between the “Passed” and “Failed” chambers when the increase in respiration exceeded 20% (15.8 pmol·s^-1^·mg^-1^ or 21.1% decrease in “Failed” chambers relative to “Passed” chambers, *P* = 0.026), but not when the increase in respiration was 10 to 15% (7.1 pmol·s^-1^·mg^-1^ or 7.9% increase in “Failed” chambers relative to “Passed” chambers, *P* = 0.563) or 15 to 20% (16.1 pmol·s^-1^·mg^-1^ or 16.1% increase in “Failed” chambers relative to “Passed” chambers, *P* = 0.387; Figure 3A). The CV and TE_M_ was calculated between the respiration rate of the “Failed” chamber and the mean respiration rate of two “Passed” chambers. There were no significant differences between the CV of the three groups (*P* = 0.073); however, there was a trend for a larger CV with a higher percentage increase in respiration after cytochrome *c* injection (Figure 3B). When respiration increased 10 to 15%, the TE_M_ was 13.0 pmol·s^-1^·mg^-1^ or 13.9%. When respiration increased 15 to 20%, the TE_M_ was 21.5 pmol·s^-1^·mg^-1^ or 19.9%; and when respiration increased above 20%, the TE_M_ was 18.5 pmol·s^-1^·mg^-1^ or 27.5%.

**Figure 3:**
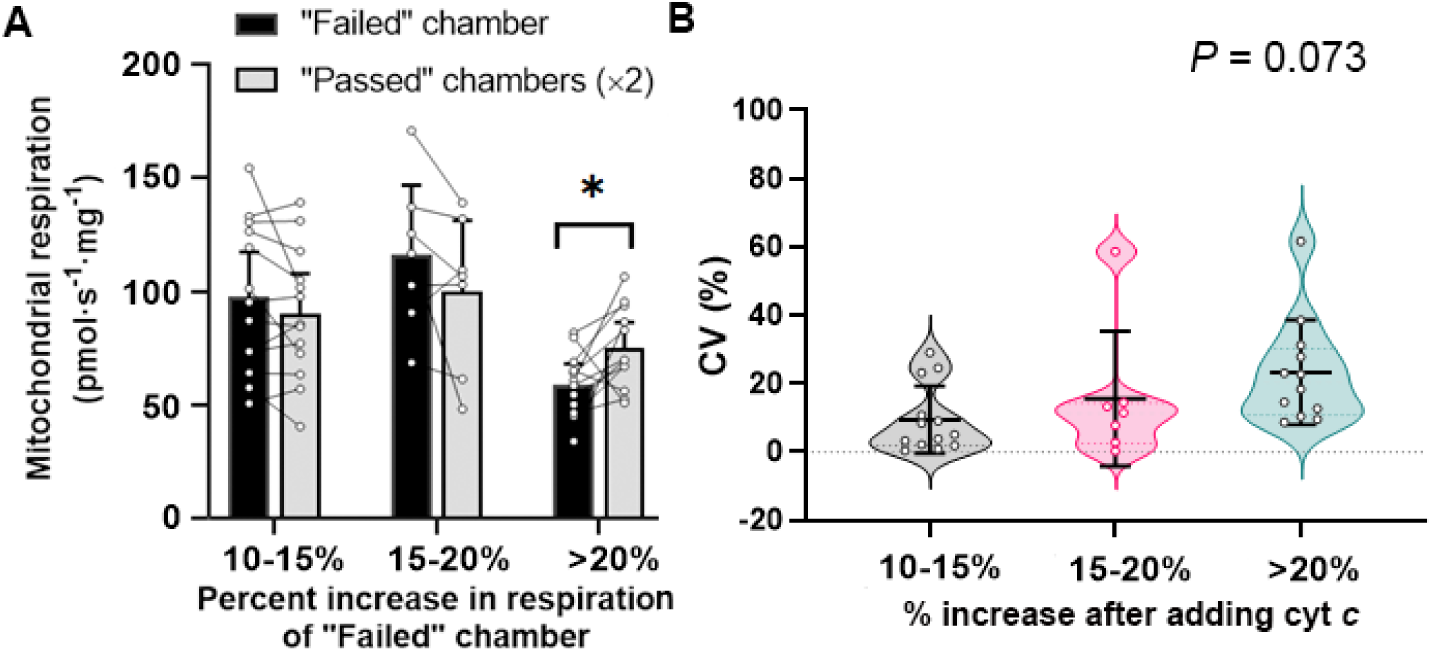
Validation of cytochrome *c* test for testing mitochondrial outer membrane integrity. **A:** Maximal mitochondrial respiration (CI + CII_p_) of permeabilized skeletal muscle (*n* = 31) measured across three chambers, where one chamber failed the cytochrome *c* test (black bars) and two chambers passed (grey bars). Data from the same assay are linked by horizontal lines. * represents *P* < 0.05. **B:** The coefficient of variation of maximal mitochondrial respiration (CI + CII_p_) between measurements from chambers failed and passed cytochrome *c* test. Individual coefficient of variation are represented by open circles; the mean and standard deviation are presented by the black line and error bars, respectively. The width of the shaded area of violin plot represents the proportion of the data located there.

### Experiment 4: Reliability of mitochondrial respiration between repeated muscle biopsies in permeabilized skeletal muscle fibre

To examine the biological variability of repeated muscle biopsies, 24 participants provided muscle samples on Day 1 and Day 6 without any intervening exercise or dietary interventions. Eight participants also provided a third muscle sample on Day 11. Each muscle sample was analyzed using three chambers across two respirometers, and all values were used for analysis without excluding any data (Supplementary Table 1 and Supplementary Table 2). However, some samples were excluded from the subsequent analysis, due to failing the outer mitochondrial membrane integrity test. There were no differences in maximal respiration (CI + CII_p_) between either two repeated muscle biopsies (*n* = 24, *mean ± SD*, Day 1: 81.8 ± 21.7 pmol·s^-1^·mg^-1^; Day 6: 80.6 ± 21.0 pmol·s^-1^·mg^-1^, *P* = 0.831; Figure 4A) or three repeated muscle biopsies (*n* = 8, *mean ± SD*, Day 1: 71.1 ± 12.9 pmol·s^-1^·mg^-1^; Day 6: 77.1 ± 21.0 pmol·s^-1^·mg^-1^; Day 11: 64.9 ± 16.0 pmol·s^-1^·mg^-1^, *P* = 0.161; Figure 4C). The average CV for two repeated respiration measurements (Day 1 and 6) was 9.7% (range from 0.9% to 46.5%; Figure 4B), and the TE_M_ was 12.2 pmol·s^-^1·mg^-1^ or 15.0%. The average CV of three repeated measurements (Day 1, 6, and 11) was 12.8% (range from 4.0% to 33.6%; Figure 4D), and the TE_M_ was 11.7 pmol·s^-^1·mg^-1^ or 16.4%. The average CV of two biological repeated measurements conducted 10 days apart (*i.e.*, Day 1 and 11) was 13.5% (range from 2.8% to 24.4%; Figure 4E), and the TE_M_ was 9.8 pmol·s^-^1·mg^-1^ or 14.4%.

**Figure 4:**
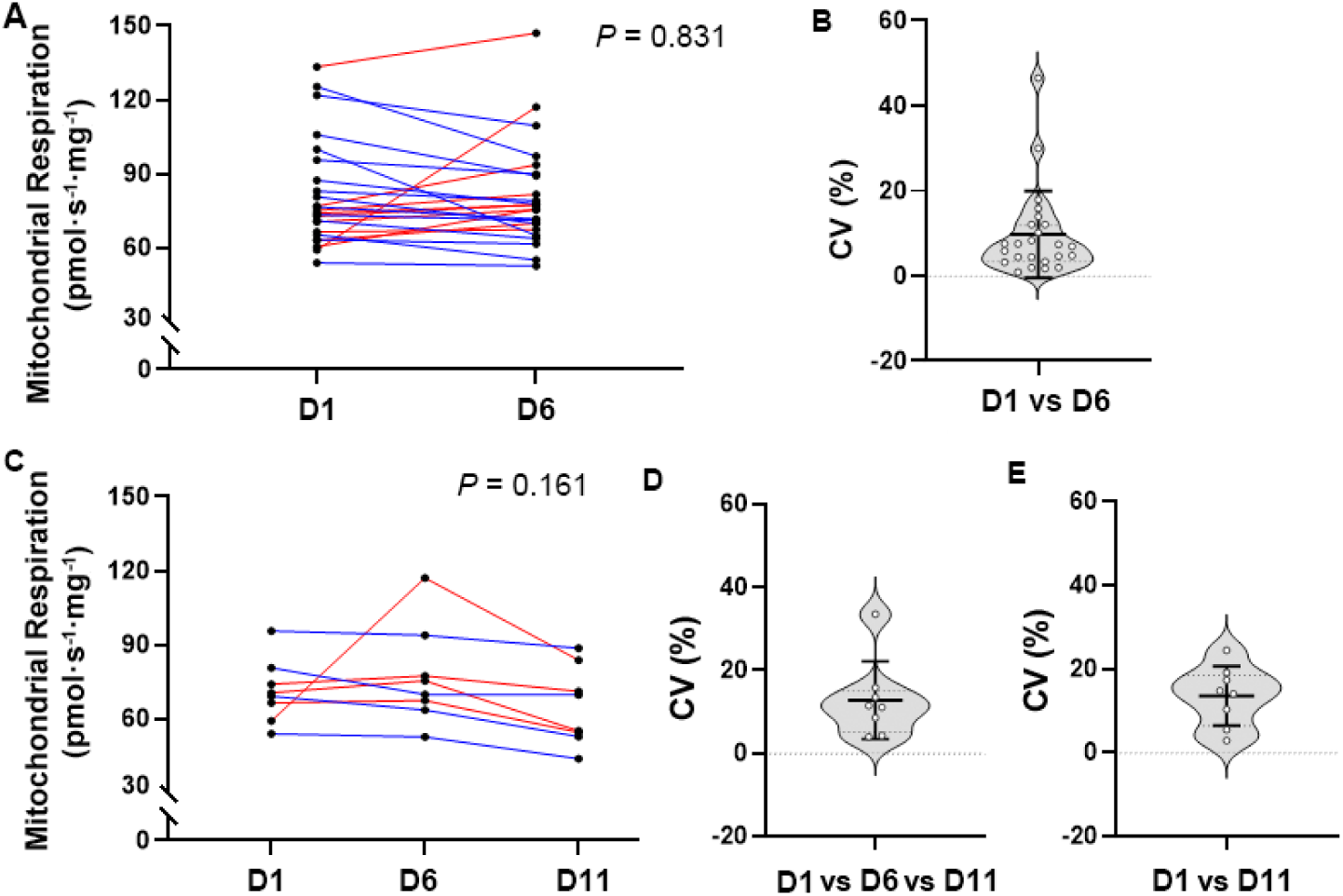
Maximal mitochondrial respiration (CI + CII_p_) of permeabilized skeletal muscle from two or three repeated muscle biopsies. **A and C:** The maximal mitochondrial respiration of (A) two repeated biopsies (*n* = 24, Day 1 and 6) and (C) three repeated biopsies (*n* = 8, Day 1, 6, and 11) are represented by black dots and connected with a line. The red lines represent an increase, whereas blue lines represent a decrease related to Day 1. **B, D, and E**: The respective coefficients of variation of (B) two maximal mitochondrial respiration measurements, 5 days apart (*n* = 24, Day 1 and 6), (D) three repeated measurements (*n* = 8, Day 1, 6, and 11), and (E) two measurements 10 days apart (*n* = 8, Day 1 and 11) are represented by open circles; the mean and standard deviation are presented by the black line and error bars, respectively. The width of the shaded area of violin plot represents the proportion of the data located there.

### Experiment 5: Reliability of CS activity between three technical repeats on the same microplate

Forty-two muscle lysates were analyzed to assess the within-plate variability between three technical repeats of CS activity determined using a microplate (*i.e.*, between three separate wells on the same plate; Supplementary Table 3). There was no significant difference between the triplicates (*P* = 0.095; Figure 5A). The average CV of the three repeated measurements was 3.5% (range from 0.2% to 14.4%; Figure 5B), and the TE_M_ was 0.11 mol·h^-1^·kg protein^-1^ or 4.1%.

**Figure 5:**
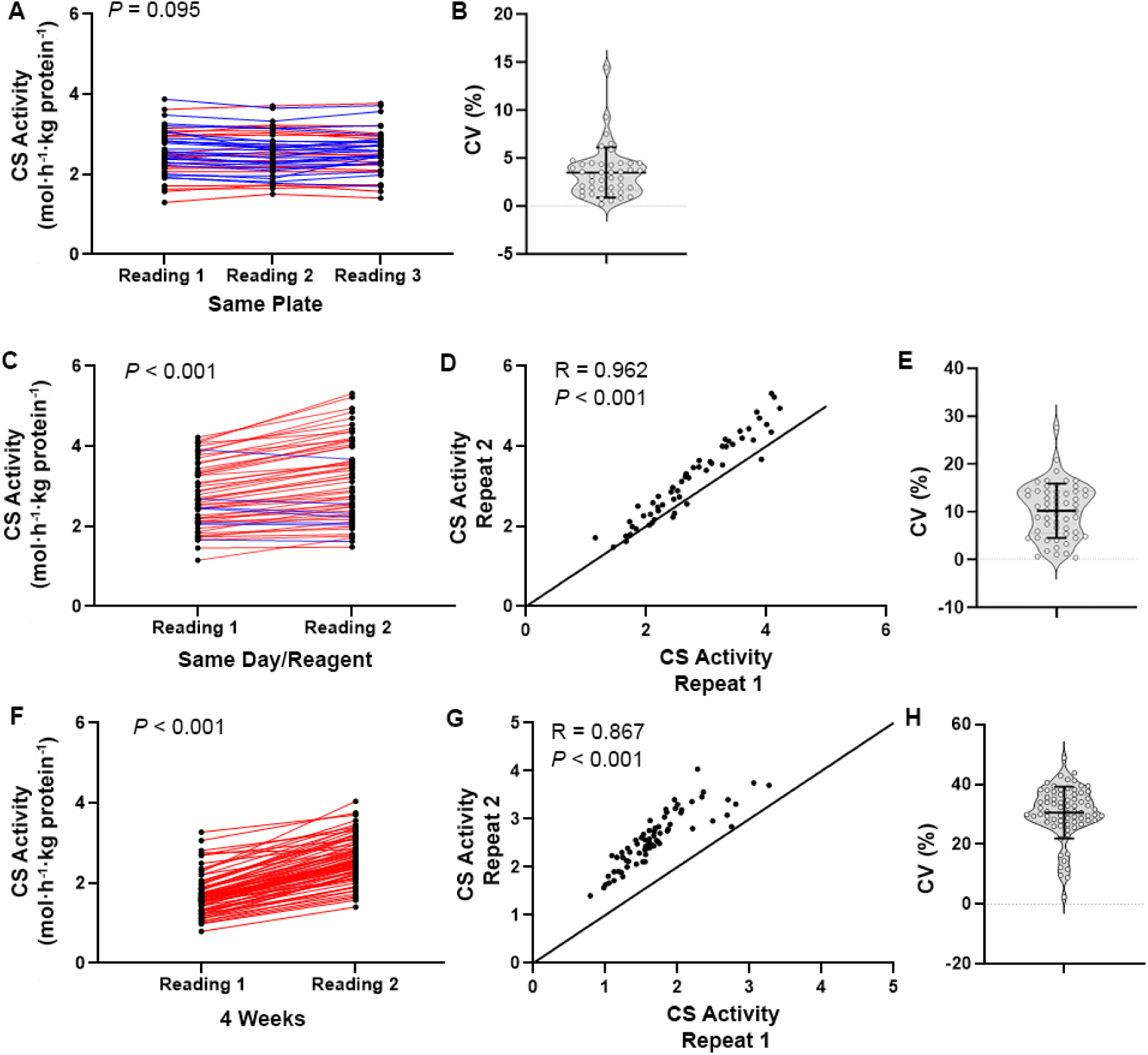
Reliability of citrate synthase activity between repeated measurements of the same skeletal muscle protein lysate. **A:** Three technical repeated measurements of citrate synthase activity (*n* = 42) determined on the same microplate are represented by black dots and connected with a line; red lines represent an increase and blue lines represent a decrease related to Reading 1. **B**: The respective coefficients of variation of the triplicate measurements are represented by open circles; the mean and standard deviation are presented by the black line and error bars, respectively. The width of the shaded area of violin plot represents the proportion of the data located there. **C and F**: Two technical repeated measurements of citrate synthase activity determined using the same skeletal muscle protein lysate in the same day (B, *n* = 55) and four weeks apart (F, *n* = 72) are represented by black dots and connected with a line; red lines represent an increase and blue lines represent a decrease between wells. **D and G**: Pearson’s correlation between the two repeated measurements in (D) the same day or (G) four weeks apart; all values are in mol·h^-1^·kg protein^-1^. The solid line shows the line of identity. **E and H**: The respective coefficients of variation of two repeated analysis in (E) the same day and (H) four weeks apart are represented by open circles; the mean and standard deviation are presented by the black line and error bars, respectively. The width of the shaded area of violin plot represents the proportion of the data located there.

### Experiment 6: Reliability of CS activity between two repeated measurements four hours apart

CS activity was measured on 55 muscle sample homogenates, twice in the same day (approximately four hours apart, with no freeze-thaw cycle in between), using the same reagents made fresh in the morning (Supplementary Table 3). The two measurements of CS activity were significantly different (0.4 mol·h^-1^·kg protein^-1^ or 15.3% increase in Reading 2 relative to Reading 1*, P* < 0.001; Figure 5C), but were correlated with each other (R = 0.962, *P* < 0.001; Figure 5D). The average CV between the two measurements was 10.2% (range from 0.4% to 27.6%; Figure 5E). The TE_M_ was 0.38 mol·h^-1^·kg protein^-1^ or 12.8%.

### Experiment 7: Reliability of CS activity between two repeated measurements four weeks apart

Using the microplate assay, CS activity was measured on 72 muscle sample homogenates, on two separate occasions four weeks apart, where sample lysates were kept at −80°C with one additional freeze-thaw cycle, and freshly made reagents were used on each occasion (Supplementary Table 3). The two measurements of CS activity were significantly different (0.9 mol·h^-1^·kg protein^-1^ or 52.7% greater in Reading 2 relative to Reading 1*, P* < 0.001; Figure 5F), but were correlated with each other (R = 0.867, *P* < 0.001; Figure 5G). The average CV between the two measurements was 30.5% (range from 2.3% to 48.8%; Figure 5H). The TE_M_ was 0.66 mol·h^-1^·kg protein^-1^ or 30.9%.

### Experiment 8: Reliability of CS activity between repeated muscle biopsies

Using the muscle samples collected in Experiment 4 (Day 1 vs Day 6 vs Day 11), CS activity was determined via the microplate assay, using freshly made reagents on the same day (Supplementary Table 2 and Supplementary Table 3). The CS activity of two repeated muscle biopsies (Day 1 vs 6), which were homogenized and measured at the same time, showed a trend to be different from each other (*n* = 24, *P* = 0.062, *mean ± SD*, Day 1: 2.7 ± 0.7 mol·h^-1^·kg^-1^; Day 6: 2.5 ± 0.4 mol·h^-1^·kg^-1^; Figure 6A). The average CV for two biologically repeated measurements (Day 1 vs Day 6) was 10.1% (range from 0.4% to 28.4%; Figure 6B), and the TE_M_ was 0.32 mol·h^-1^·kg protein^-1^ or 12.3%. The CS activity of the three repeated muscle biopsies were not significantly different from each other (*n* = 8, *P* = 0.948, *mean ± SD*, Day 1: 2.4 ± 0.6 mol·h^-1^·kg^-1^; Day 6: 2.4 ± 0.4 mol·h^-1^·kg^-1^; Day 11: 2.3 ± 0.3 mol·h^-1^·kg^-1^; Figure 6C). The average CV for three biological repeated measurements (Day 1, 6, and 11) was 12.6% (range from 2.7% to 32.5%; Figure 6D), and the TE_M_ was 0.4 mol·h^-1^·kg protein^-1^ or 15.6%. The average CV for two biological repeated measurements 10 days apart (*i.e.*, Day 1 and 11) as 14.1% (range from 0.4% to 40.8%; Figure 6E), and the TE_M_ was 0.43 mol·h^-1^·kg protein^-1^ or 18.5%.

**Figure 6:**
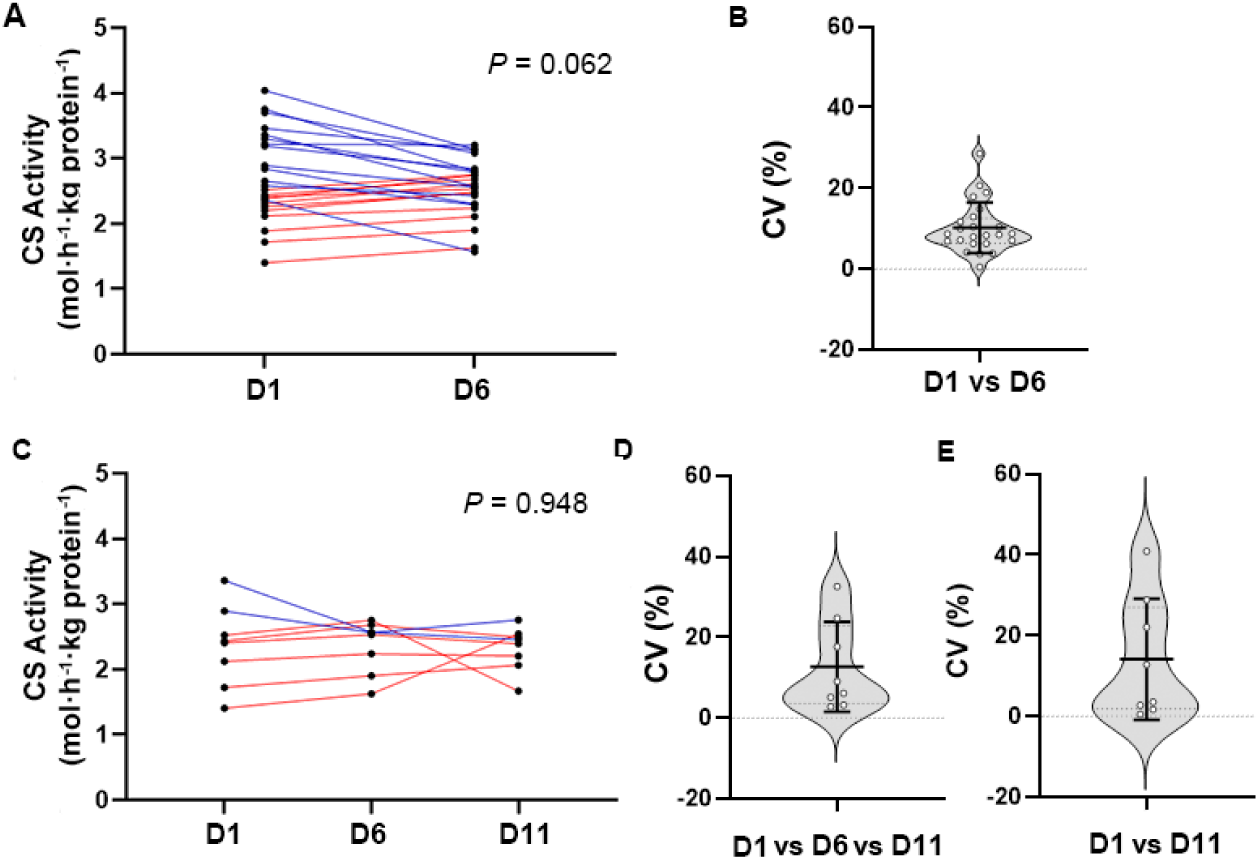
Citrate synthase activity, determined via microplate assay using repeated muscle biopsies. **A and C:** Citrate synthase activity of two repeated biopsies (Day 1 and 6, *n* = 24) and three repeated biopsies (Day 1, 6, and 11, *n* = 8) are represented by black dots and connected with a line. The red lines represent an increase, whereas blue lines represent a decrease related to Day 1. **B, D, and E**: The respective coefficients of variation of (B) two repeated citrate synthase activity measurements (Day 1 and 6, *n* = 24), (D) three repeated citrate synthase activity measurements (Day 1, 6, and 11, *n* = 8), and (E) repeated citrate synthase activity measurements 10 days apart (Day 1 and 11, *n* = 8) are represented by open circles; the mean and standard deviation are presented by the black line and error bars, respectively. The width of the shaded area of violin plot represents the proportion of the data located there.

### Experiment 9: Reliability of CS activity between two measurements using different protocols (microplate and cuvette)

In addition to the microplate assay, CS activity can also be measured in cuvettes. To determine if different measurement techniques/protocols influences CS activity measurements, we compared CS activity using both protocols. The CS activity of 40 skeletal muscle samples was determined via both microplate and cuvette protocols, in two independent laboratories (Supplementary Table 3). CS activity of the two protocols were significantly different (1.3 mol·h^-1^·kg protein^-1^ or 19.4% lower in cuvette relative to microplate, *P* < 0.001; Figure 7A), but correlated (R = 0.707, *P* < 0.001, (Figure 7B). The average CV between the two measurements was 17.0% (range from 0.7% to 48.7%; Figure 7C). The TE_M_ was 1.3 mol·h^-1^·kg protein^-1^ or 21.9%. The Lin’s concordance correlation coefficient (p_c_) was 0.472 (95% confidence interval, 0.308 to 0.635). The average difference (bias) between the two methods was 1.275 mol·h^-1^·kg protein^-1^ (SD of bias = 1.340, Bland-Altman plot; Figure 7D).

**Figure 7:**
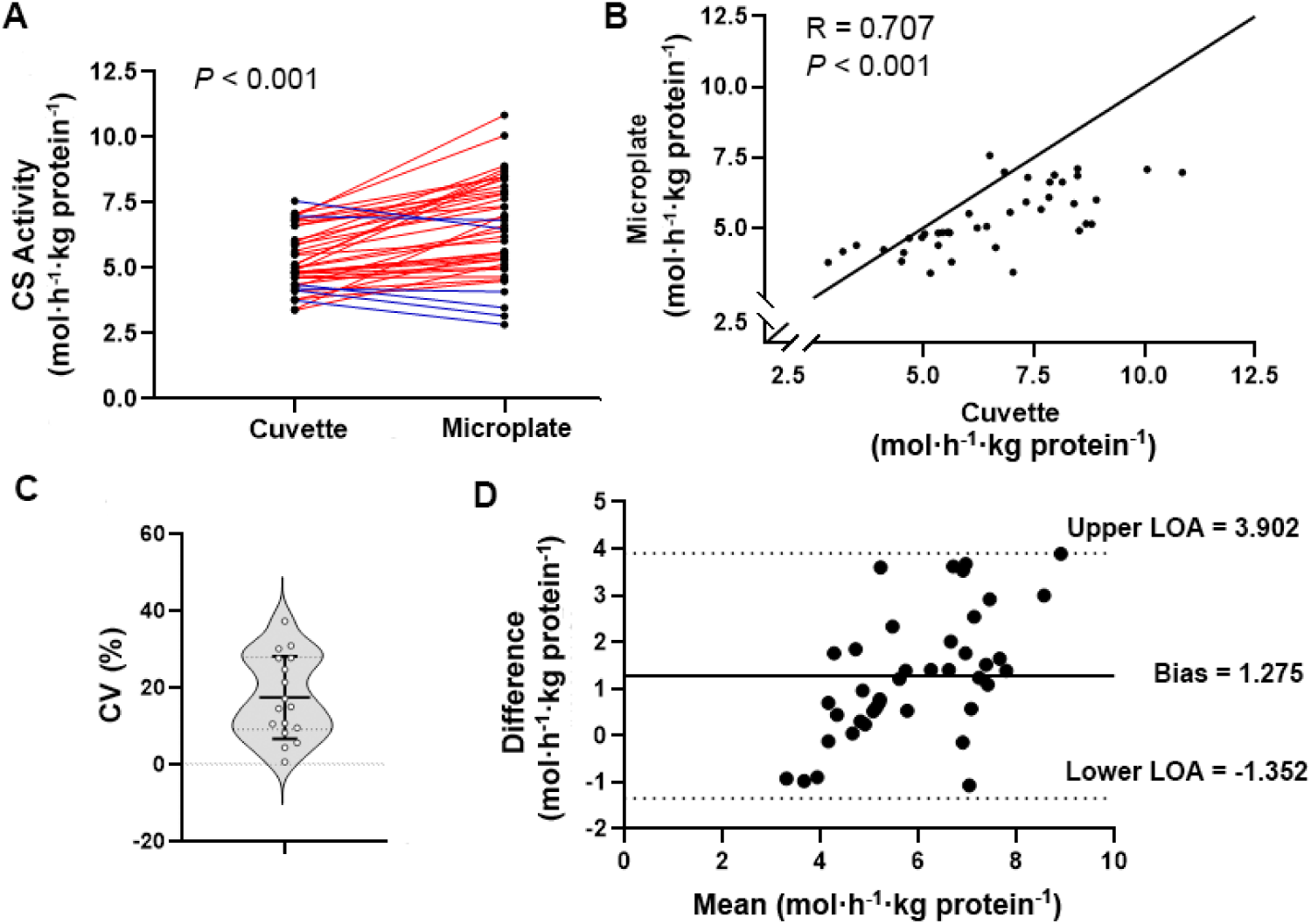
**A:** Citrate synthase (CS) activity of skeletal muscle (*n* = 40) derived using two different protocols (cuvette vs microplate) are represented by black dots and connected with a line; red lines represent an increase, and blue lines represent a decrease with the microplate reading. **B:** Pearson’s correlation between the two measurements; all values are in mol·h^-1^·kg protein^-1^. The solid line shows the line of identity. **C**: The respective coefficients of variation between the two protocols are represented by open circles; the mean and standard deviation are presented by the black line and error bars, respectively. The width of the shaded area of violin plot represents the proportion of the data located there. **D**: Bland-Altman plot showing the agreement between the two methods for determining CS activity. The difference in CS activity between protocols is shown on the y-axis, the mean CS activity of the two protocols is shown on the x-axis (both in mol·h^-1^·kg protein^-1^). The solid line represents the mean difference between the two protocols (*i.e.*, bias). The two dashed lines represent the Limits of Agreement (LOA = 1.96 x SD of the mean difference between the two methods).

### Experiment 10: Reliability of mitochondrial specific respiration between repeated muscle biopsies

Mitochondrial-specific respiration is often reported as mass-specific respiration normalized to mitochondrial content. Here, we have calculated mitochondrial-specific respiration using the data obtained in Experiments 4 and 7 (Supplementary Table 2 and Supplementary Table 4). Mitochondrial-specific respiration did not differ significantly between two repeated muscle biopsies (*n* = 24, Day 1 and 6, *mean ± SD*, Day 1: 30.2 ± 6.2 mol·h^-1^·CS^-1^; Day 6: 30.2 ± 6.8 mol·h^-1^·CS^-1^, *P* = 0.266) nor three repeated muscle biopsies (*n* = 8, Day 1, 6, and 11, *mean ± SD*, Day 1: 31.4 ± 7.0 mol·h^-1^·CS^-1^; Day 6: 33.0 ± 7.4 mol·h^-1^·CS^-1^; Day 11: 28.1 ± 6.0 mol·h^-1^·CS^-1^, *P* = 0.397), respectively (Figures 8A and C). The average CV for two repeated measurements (Day 1 and 6) was 12.5% (range from 0.2% to 54.0%; Figure 8B), and the TE_M_ was 5.5 mol·h^-1^·CS^-1^ or 17.6%. The average CV for three repeated measurements (Day 1, 6, and 11) was 19% (range from 1.8% to 39.5%; Figure 8D), and the TE_M_ was 6.9 mol·h^-1^·CS^-1^ or 22.4%. The average CV of two repeated measurements 10 days apart (*i.e.*, Day 1 and 11) was 21.1% (range from 2.4% to 54.0%; Figure 8E), and the TE_M_ was 7.5 mol·h^-1^·CS^-1^ or 25.1%. In addition, we determined that mass-specific maximal respiration rate (CI + CII_p_) from Experiment 4 was correlated with CS activity from Experiment 7 (*n* =56, R = 0.587, *P* < 0.001; Figure 8F).

**Figure 8:**
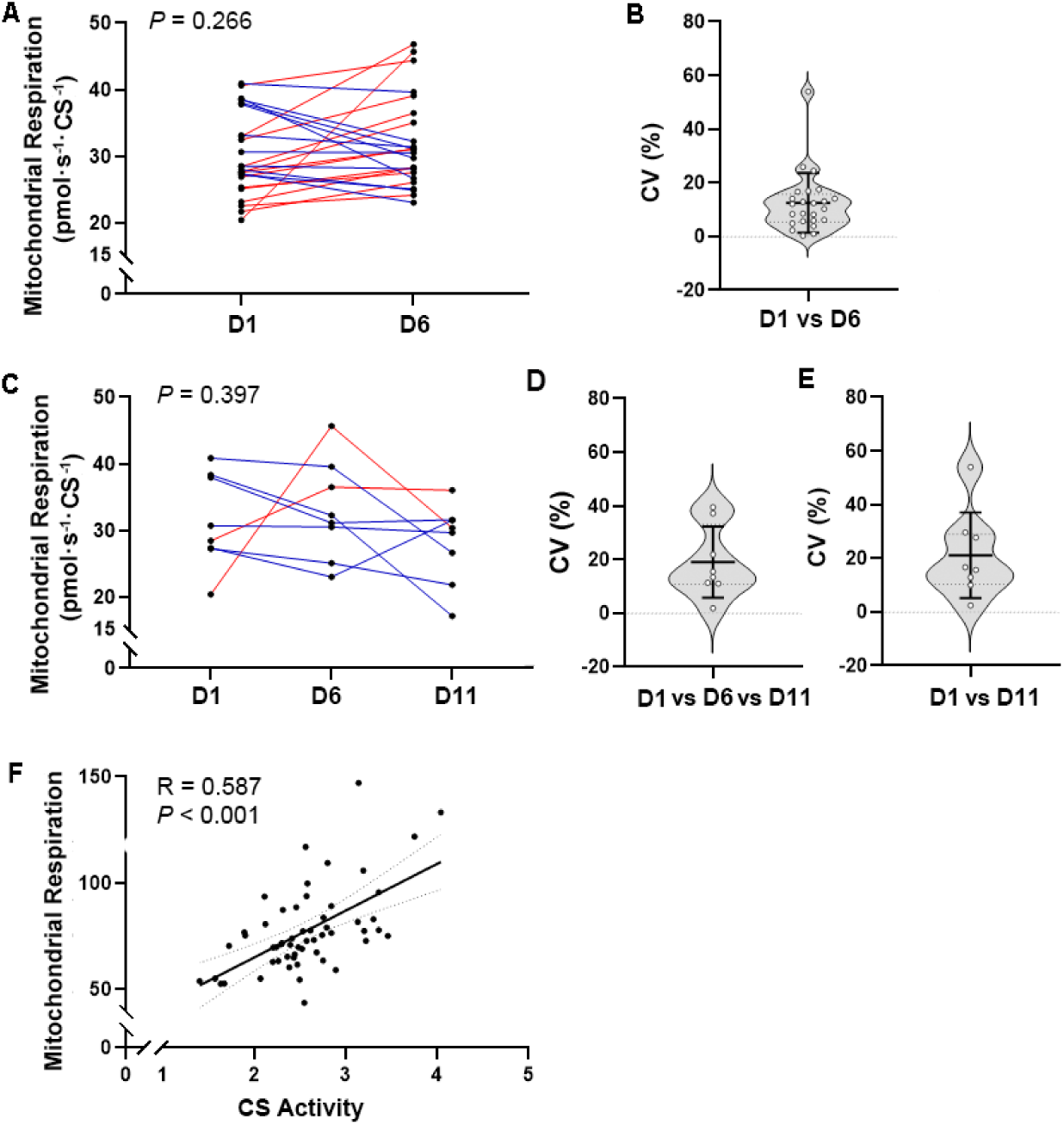
Mitochondrial-specific respiration using repeated muscle biopsies. **A and C**: Mitochondrial-specific respiration of (A) two repeated biopsies (Day 1 and 6, *n* = 24) and (C) three repeated biopsies (Day 0, 5 and 10, *n* = 8) are represented by black dots and connected with a line. The red lines represent an increase, whereas blue lines represent a decrease related to Day 1. **B, D, and E:** The respective coefficients of variation for (B) two measurements (Day 1 and 6, *n* = 24), (D) three measurements (Day 1, 6 and 11, *n* = 8), and (E) two measurements 10 day apart (Day 1 and 11, *n* = 8) are represented by open circles; the mean and standard deviation are presented by the black line and error bars, respectively. The width of the shaded area of violin plot represents the proportion of the data located there. **F:** Pearson’s correlation between citrate synthase activity (mol·h^-1^·kg protein^-1^) and maximal mitochondrial respiration (pmol·s^-1^·mg^-1^), using data obtained in Experiments 4 and 7 (n = 56). Simple linear regression (solid line) with 95% confidence internals (dash line) is shown.

### Power analysis for detecting meaningful changes in mitochondrial respiration and content

It is possible to calculate the power to determine changes in mitochondrial respiration and mitochondrial content with different sample sizes based on the mean result, standard deviation (SD), and absolute TE_M_ obtained from each experiment [19]. The minimal change in maximal mitochondrial respiration that can be detected with 80% power with 10 and 20 participants is 17% and 11% for Experiment 2 (two repeated measurements; Figure 9A), 20% and 13% for Experiment 3 when using all three repeated measurements (“All Three Values” results from three repeated measurements; Figure 9A), 13% and 9% when outliers were excluded from the calculation (“Values Within 4× CV” results from three repeated measurements; Figure 9A), 18% and 12% for Experiment 4 (three repeated muscle biopsies in Day 1, 6, and 11; Figure 9B). We also determined the minimum detectable changes in CS activity. For Experiment 5 (three repeated measurements on the same plate), the minimum change in CS activity that can be detected with 80% power with 10 and 20 participants is 5% and 3% (Figure 9C). For Experiment 8 and 9 (three repeated muscle biopsies in Day 1, 6, and 11), the minimum change in CS activity (Figure 9D) that can be detected with 80% power with 10 and 20 participants is 18% and 12% respectively; the minimum change in mitochondrial-specific respiration (Figure 9E) that can be detected with 80% power with 10 and 20 participants is 24% and 16% respectively.

**Figure 9:**
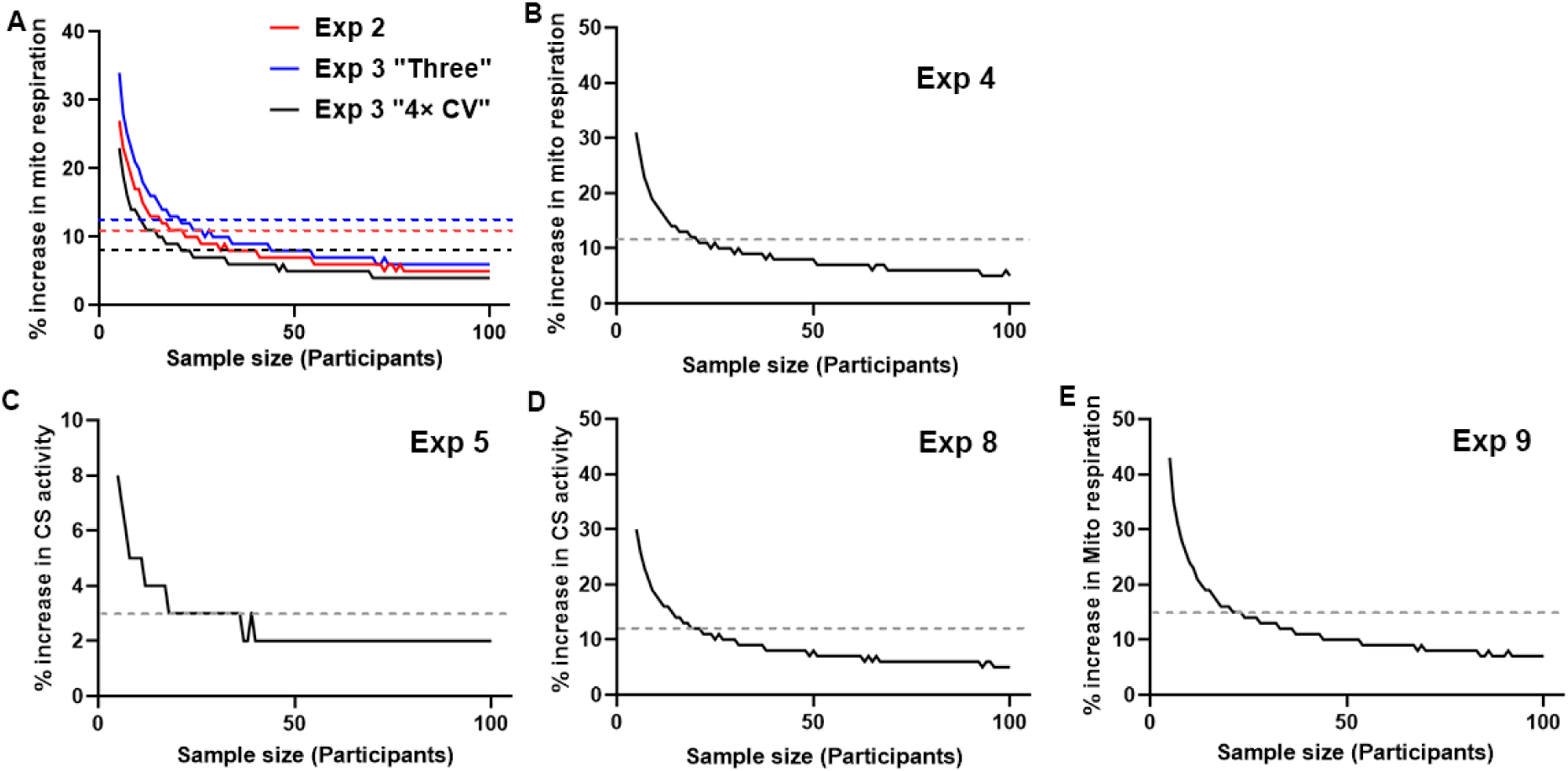
Simulations to estimate the sample size required to detect changes in maximal respiration (CI + CII_p_) and mitochondrial content with 80% power. **A and B:** The minimal sample size required for (A) Experiment 2 (red line) and 3 (blue and black lines) and (B) Experiment 4 to detect an increase in maximal mitochondrial respiration after an intervention with 80% power. **C and D:** The minimal sample size required for (C) Experiment 5 and (D) Experiment 8 to detect an increase in CS activity after intervention, with 80% power. **E:** The minimal sample size required for Experiment 9 to detect an increase in mitochondrial-specific respiration after intervention, with 80% power. Vertical dash line indicates changes can be detected with sample size of 20.

## DISCUSSION

Mitochondrial adaptations to exercise training are of great interest to optimise training prescriptions and health. However, measurements of both mitochondrial respiration and mitochondrial content are affected by variations in the protocols employed by individual research laboratories, which in turn can influence the findings of individual studies. We have investigated the reliability of measurements of mitochondrial respiration and mitochondrial content in human skeletal muscle. We provide new data, including the variability of technical and biological repeated measurements, the variability between different protocols, and conducted power analysis to determine the minimal changes that can be detected in experiments. We will discuss how these methodological considerations might affect the design, execution, and interpretation of the results of future studies.

### Variability in mitochondrial respiration between technical replicates

Many factors contribute to the variation between technical repeated measurements of mitochondrial respiration (*i.e.*, analyzing muscle fibres from the same biopsy sample in two chambers), including the way muscle biopsies are obtained, the degree of mechanical separation of muscle fibres from each portion of muscle, the accuracy in measuring the muscle mass, as well as the respirometer itself. To assess the analytical variation caused by the imperfections in experimental methods and instruments, we measured the respiration rate using a yeast cell suspension and report a CV of 6.1%, which is consistent with a CV of 5.4% reported elsewhere [33]. This indicates that approximately 6% of the variability in measurements stem from the background or “noise” from the machine, and operator inaccuracy. However, it is important to underline that the level of background or “noise” depends on factors such as the maintenance and calibration of the machines; therefore, the 6% CV value we report is the background of respirometers from our laboratory.

When analyzing the same muscle sample in duplicate, we identified a greater CV (12.7%) and TE_M_ (12.8 pmol·s^-1^·mg^-1^ or 15.4%); however, our observed variability is in a similar range to, or slightly lower than, previous studies [19, 32, 33]. Sahl *et al.* reported a CV of 8.4% [33], and Cardinale *et al.* [32] reported a CV of 15.2% and TE_M_ of 10.5 pmol·s^-1^·mg^-1^ for duplicate analyses on the same muscle sample. Jacques *et al.* [19] also reported that a TE_M_ for maximal mitochondrial respiration (CI+CII_p_) of 19 pmol·s^-1^·mg^-1^ or 15.3%. The comparable variability between the present study and existing literature suggests that the measurement variation between replicates is due to both the machine itself (as shown by the yeast cell suspension, where there is no biological variation in the samples assessed in the two separate chambers) as well as differences in sample handling and biological variability (evident from the increased CV in the muscle samples). Thus, whilst proper maintenance and calibration of the respirometer are important, standardizing sample preparation across a study, and potentially between studies, is also critically important. However, it is clear that there is substantial biological variation when measuring mitochondrial respiration in permeabilized muscle fibres using high-resolution respirometry, which in turn can interfere with reporting statistically significant differences before and after experimental intervention. Using a previously established method of statistical simulation to determine appropriate sample size [19], we calculated that two technical repeats of skeletal muscle samples from 20 participants are able to detect differences in maximal mitochondrial respiration of greater than 11% with 80% power, and a typical human experiment with 10 participants are able to detect differences in maximal mitochondrial respiration of greater than 17% with 80% power.

The relatively large variability led us to consider ways in which this variability could be minimised. The majority of studies assay respiratory function using two technical repeats (i.e., two chambers of one machine). In our laboratory, using multiple respirometers, we were able to examine the influence of using three technical repeats (*i.e.*, three chambers, across two machines). By setting the threshold for a outlier as four times the CV of yeast respiration (“machine background”) from the mean of the three technical repeats and excluding the outliers, we found that the maximal respiration calculated (“Values Within 4× CV”) was not significantly different from when all values from three chambers were used for calculation. However, the CV and TEM for the “Values Within 4× CV” were greatly reduced compared to the “All Three Values” (CV: 11.0% (“Values Within 4× CV”) vs 16.7% (“All Three Values”); TEM: 11.7% (“Values Within 4× CV”) vs 18.1% (“All Three Values”)). Using statistical simulations [19], we then calculated that 10 and 20 samples measured in triplicate (using all values for calculation) could determine 20% and 13% difference in maximal mitochondrial respiration with 80% power, respectively. After excluding outliers (“Values Within 4× CV”), we then calculated that sample size of 10 and 20 measures would be able to determine 13% and 9% difference in maximal mitochondrial respiration with 80% power. In human studies looking for small differences in mitochondrial respiration and where numbers of participants are limited, this approach (excluding outliers) can reduce the technical variability and increase the power to detect smaller changes, without influencing the results (the maximal respiration rate remained the same in this present study). However, there are no strict statistical rules for identifying outliers, especially with a small sample size (*i.e.*, three repeated measurements from three chambers); thus, the threshold for defining outliers needs to be selected carefully and conservatively based on the background or “noise” level of the respirometer, as well as the study design (*i.e.*, sample size, changes in respiration expected). In this present study, a relatively large threshold was used to only remove values that were much higher or lower compared to the rest (likely to be a “true outlier”), whilst keeping values that may differ due to the inherent variability of human skeletal muscle fibres. We recommend conducting yeast respiration experiments as a routine test to monitor the background or “noise” of the respirometers and to appropriately categorise outliers. The same criteria for defining outlier needs to be applied to all data from the same study (*i.e.*, before and after an experimental intervention).

### Variability in mitochondrial respiration between biological replicates

In addition to technical variability, it is crucial to consider the biological variation when assessing changes in mitochondrial respiration, as well as the variability caused by preparation techniques and the muscle biopsy procedure *per se*. Our observed CV ranged between 9.7 to 13.5% when two or three separate muscle biopsies from the same participants were repeated over a 5- to 10-day control period. This CV range is similar to a previously reported CV of 13.3% for three repeated muscle biopsies sampled within 20 min [33], but much lower than another reported CV of 33.1% for two repeated muscle biopsies (sampled 27 ± 6 days apart) [32]. By performing statistical simulations based on the mean respiration, SD, and TE_M_ values from three repeated biopsies of 8 participants, which include both technical and biological variation, we identified that training studies with 10 and 20 participants would be able to determine 18% and 12% change in maximal mitochondrial respiration with 80% power, respectively.

### Influence of muscle sample size used for respiration measurements

Another potential cause of measurement variability may be the muscle sample mass (mg). In the present study, we found no differences in variability between technical repeats when larger (> 3 mg) or smaller (< 3 mg) muscle samples were used in both chambers. Although it is advised that 1 to 2 mg wet weight of skeletal muscle is sufficient for one chamber of Oxygraph-2k respirometer [44], we recommend to use 2-3 mg of muscle fibres per chamber, as it is easier to handle during weighing and transfer. It is also important to obtain an accurate muscle mass value, as the respiration rate is measured in relation to muscle mass. We recommend using a balance with readability range between 0.001 mg and 0.01 mg, and calibrating the balance regularly.

### Influence of absolute mitochondrial respiration value

Our data also indicate that the absolute value of mitochondrial respiration could impact the technical reliability in mitochondrial respiration. In both Experiments 2 and 3, the absolute TE_M_ values were similar, regardless of the absolute respiration rate. However, the CV and TE_M_ (expressed as a %) decreased with an increase in absolute respiration rate, as those measurements were calculated relative to the average respiration rate of all samples. As a result, we recommend taking the absolute mitochondrial respiration value of the participants into consideration during the study design process. For example, less relative variation is expected among trained participants who have greater rates of mitochondrial respiration. In contrast, older participants [45], or other clinical populations [46], for whom lower mitochondrial respiration rates are expected, may elicit more substantial variation, and therefore a larger number of participants or technical replicates might be required to achieve adequate power.

### Cytochrome *c* test for outer mitochondrial membrane integrity

Both the muscle biopsy and membrane permeabilization processes can damage the mitochondrial outer membrane and release cytochrome *c*, which is an electron-carrying protein loosely bound to the inner mitochondrial membrane [47]. When the mitochondrial outer membrane is damaged, cytochrome *c* leaks out and the respiratory capacity becomes reduced; therefore, an increase in respiration observed after adding cytochrome *c* reflects the leakiness of the outer membrane [47]. The compromised mitochondrial membrane integrity can cause misleading results and thus should be excluded from the data analysis. A common practice (also adopted in our experiments) is to accept up to a 10% increase in respiration rate following the addition of cytochrome *c* [43]; however, this acceptable threshold has not been properly validated, and some research suggests that a 10 to 20% increase in respiration could be acceptable [48]. Based on our results, the maximal respiration rate was significantly reduced only when the increase in respiration rate observed after adding cytochrome *c* exceeded 20%. However, due to the greater variability observed when the increase in respiration rate was between 15 to 20% (Figure 7B), we suggest using a threshold of 15% as the acceptable increase in mitochondrial respiration rate following the cytochrome *c* addition.

### Variability in CS activity between technical replicates

Our CV observed from three technical repeats of CS activity on the same microplate (3.5%) is consistent with other research groups (< 5%, [37]), as well as the intra-assay CV reported for a commercial human CS activity assay kit (4.4 to 6.6%, [49]). This relatively low level of variation between technical repeats is below the typical significant changes in CS activity observed following cycling training interventions (ranging from 10 days to 10 weeks), which have been reported to be 9 to 75% [5]. When performing the statistical simulation using our CS activity data, we determined that studies with 10 and 20 participants can identify 5% and 3% difference in CS activity at 80% power, respectively.

When the CS activity assay was conducted twice on the same day (four hours apart), with the same reagents, using the same muscle homogenate (kept on ice in between assays), the variability between the two measurements was greater than the within-plate variation (CV of 10.2%). However, when the CS activity assay was conducted on two separate days (four weeks apart) using the same muscle homogenate (kept at −80°C in between assays), with freshly made reagents on each occasion, the variability between the two measurements increased dramatically (CV of 30.5%), to within the typical range of changes in CS activity reported to occur with training (9 to 75%) [5]. Our observed CV from the same day is similar to the inter-assay variation reported for a commercial human CS activity assay kit (8.3% between 3 days) [49]. Nevertheless, majority of the samples analyzed four hours later and all samples analyzed four weeks later exhibited greater CS activity readings than the first assay. Given the time frame, and that the two assays were carried out by the same experienced researcher, the variation is unlikely due to protein degradation or human error, but due to the imperfections in the experimental method. One possibility is that our samples were not sonicated, thus the mitochondrial membrane was not fully disrupted and CS activity was underestimated in the earlier measurement. We recommend homogenizing all samples on the same day, and employing a sonication step that might help to reduce the technical variability. We also recommend conducting CS activity assays for all samples at the same time using the same reagents, and that samples that will be compared directly (*e.g.*, samples taken from the same participant, before and after an intervention) should be measured on the same microplate. However, if this is not feasible for some study designs, the researcher should consider including an internal control to correct the variation between assays, and consider using commercial kits that exhibit less inter-assay variation.

### Variability in CS activity between biological replicates

As with mitochondrial respiration, it is important to consider the impact of biological variation when assessing change in CS activity. Despite the different levels of technical variability observed between mitochondrial respirometry and the CS activity assay, both methods exhibited similar CV between biological repeats (Respiration: 10.1-14.1%; CS activity: 9.7-13.5%). As with our mitochondrial respirometry data, by performing statistical simulations on the CS activity data from three repeated biopsies of 8 participants, studies with 10 and 20 participants would be able to determine 18% and 12% change in CS activity with 80% power, respectively.

### Variability in CS activity using two protocols

There are considerable differences between research laboratories in the protocols and techniques used to determine CS activity. Once such example is the use of a microplate or a cuvette when measuring CS activity using a spectrophotometer. In the present study, a portion of the muscle biopsy was homogenized in our laboratory and measured with a microplate protocol. Another portion of the muscle biopsy was sent to a collaborator’s laboratory, where muscle was homogenized using a separate protocol and CS activity was measured using cuvette. The large variability and the difference in CS activity between two measurements (1.28 ± 1.34 mol·h^-1^·kg protein^-1^) relative to the mean of the two methods (cuvette = 5.31 mol·h^-1^·kg protein^-1^, microplate = 6.58 mol·h^-1^·kg protein^-1^) suggest that the results from the two protocols should not be compared directly. The Lin’s concordance correlation coefficient (p_c_) for two different protocols was also considered poor [50]. As a result, the CS activity from different studies should not be directly compared if the protocols used are not standardised. Instead, the relative change in CS activity after the intervention could provide a more appropriate means to compare and interpret results between studies and protocols [5].

### Variability in mitochondrial-specific respiration between biological replicates and correlation between mass-specific respiration and CS activity

Mitochondrial-specific respiration, whereby mass-specific respiration obtained by high-resolution respirometry is corrected for any changes in mitochondrial content, can provide additional information on mitochondrial function (*i.e.*, whether the change in mitochondrial respiration is due to changes in intrinsic mitochondrial characteristics or whether it stems from changes in mitochondrial content). Thus, it has become commonplace to analyze mitochondrial respiration relative to mitochondrial content or one of its valid biomarkers to elucidate intrinsic mitochondrial adaptations [9, 39, 51]. The variability in mitochondrial-specific respiration (Experiment 9, 12.5-21.1%) is greater than that in mass-specific respiration (Experiment 4, 9.7-13.5%), which could be due to a combination of technical variability from the measurements of both mitochondrial respiration and mitochondrial content. Statistical simulations on data from three repeated biopsies of 8 participants determined that studies with 10 and 20 participants would enable to determine 24% and 16% change in mitochondrial-specific respiration with 80% power (compared to a 12% change in mass-specific respiration and CS activity).

## Conclusion

Based on our findings, we have provided some important technical considerations for measuring mitochondrial respiration in human skeletal muscle. Firstly, the variability in mitochondrial respiration is considerably high when muscle samples are analyzed in duplicate. The technical variability can be reduced if the assay is performed in triplicate (using three chambers), and only values within the acceptable range, which should be defined by the individual study, are used for data analysis. By reducing the technical variability, the power to detect smaller changes after an experimental intervention will be increased, meaning a smaller sample size will be required for the study design. Secondly, the biological variability and absolute mitochondrial respiration value of the participants should be considered when estimating the required sample size. Studies aiming to detect small changes in mitochondrial respiration may require two baseline measurements over a control period. Thirdly, we suggest using 15% as the threshold for the increase in respiration rate after the addition of cytochrome *c* test for testing mitochondrial outer membrane integrity.

For CS activity, a biomarker of mitochondrial content, there are several technical considerations required when designing and performing the assay. Firstly, all samples from the same study should be homogenized at the same time, then measured at the same time using the same batch of freshly made chemical reagents. Secondly, similar to measurements of mitochondrial respiration, biological variability, which is greater than the technical variability for the CS activity assay, should be considered, if a small change in mitochondrial content is expected. Finally, the relative change in CS activity, rather than the absolute value, should be used to compare the outcomes from different studies using non-standardized protocols. By implementing methodologies incorporating these technical considerations, the variability within these mitochondrial measures of human skeletal muscle can be minimised and give greater confidence when interpreting results.

## METHODS

### Ethics approval

All experiments were approved by the Victoria University Human Research Ethics Committee. Participants were given comprehensive, written and verbal information about the original experiments, before providing written consent to participate, and have their data used in this study.

### Study design

The study design for each experiment is summarised in Figure 10. All human skeletal muscle samples were obtained at rest (*i.e.*, no exercise was performed in the 48 h prior).

**Figure 10:**
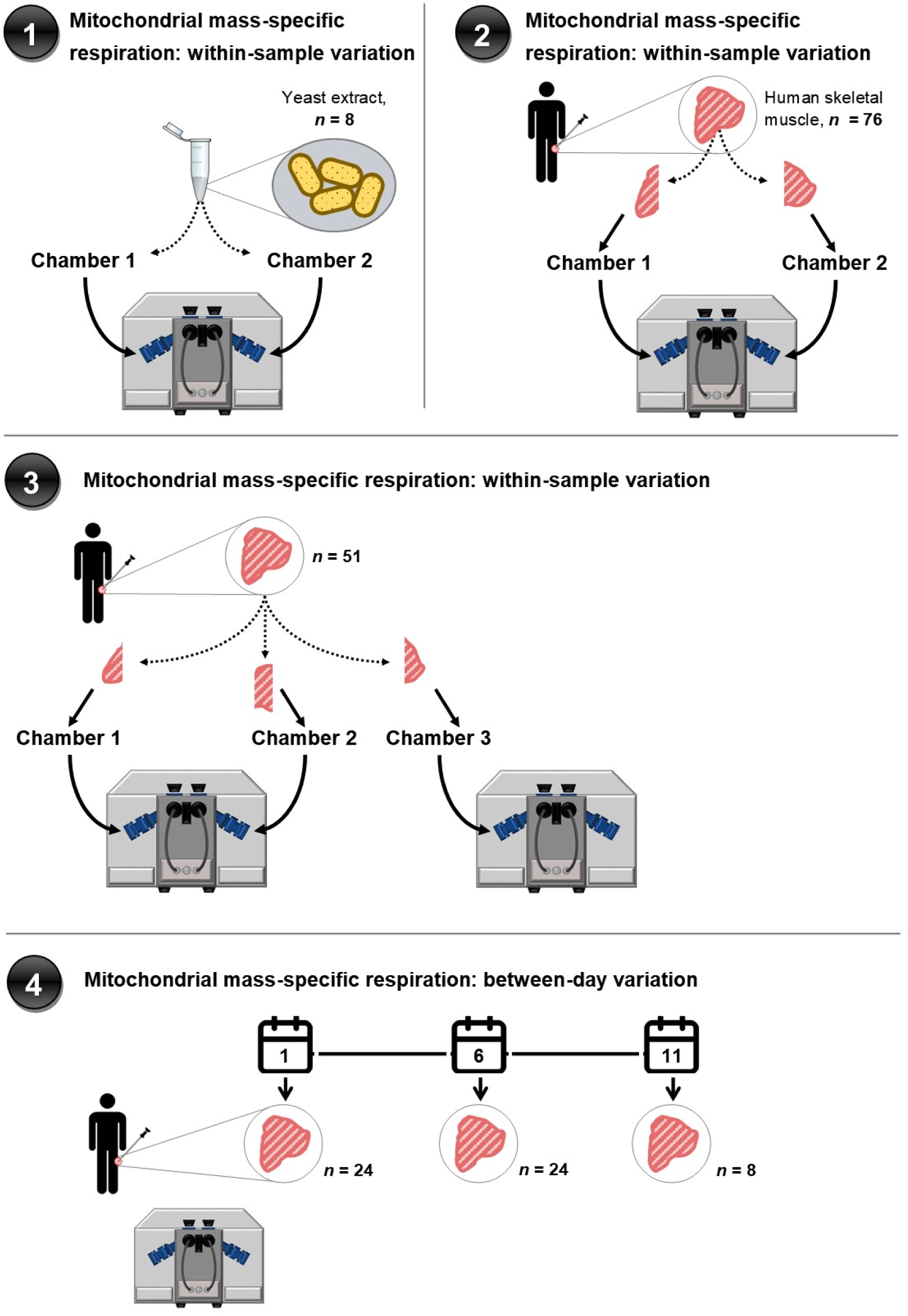
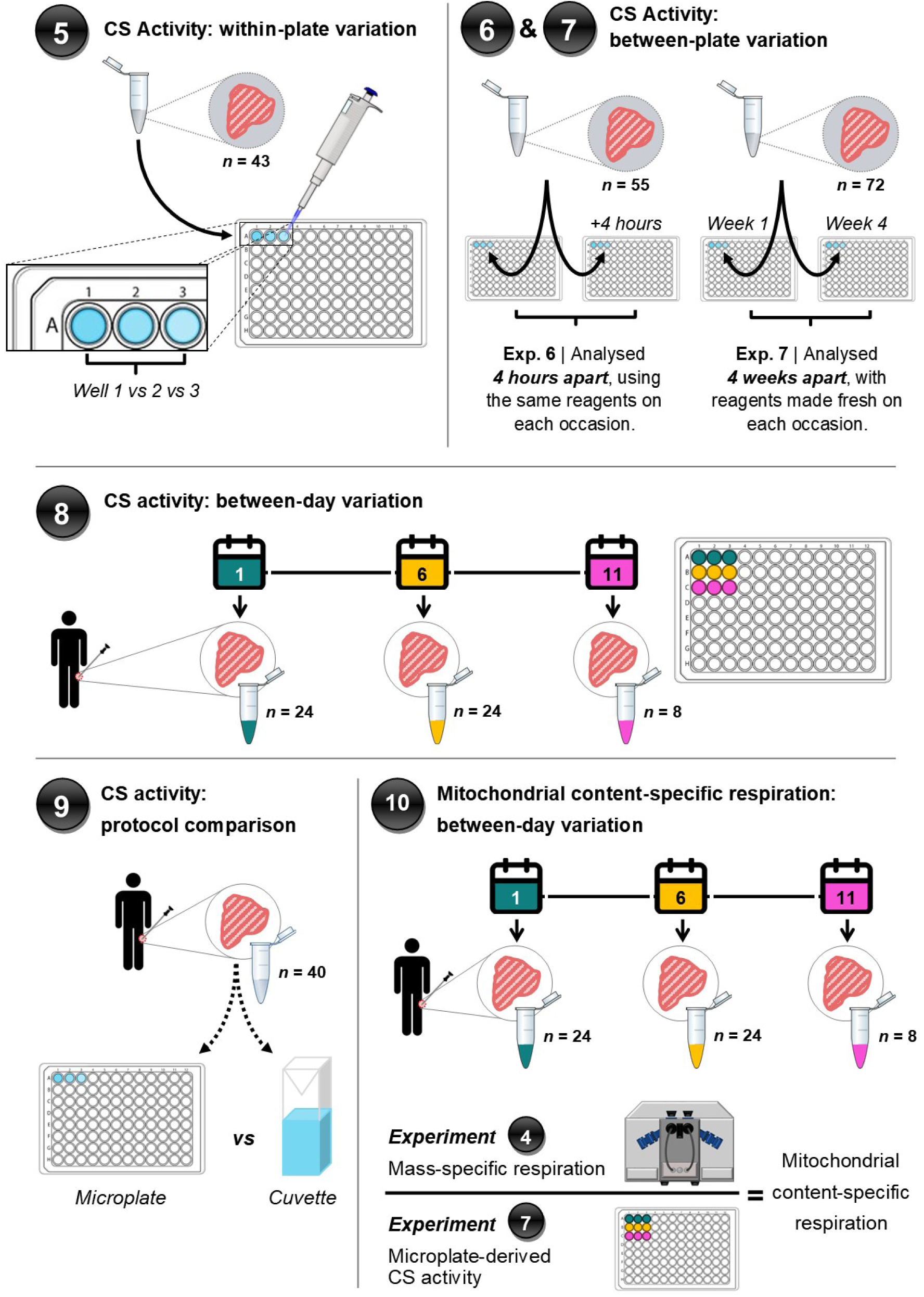
Outline of the study design. Nine experiments were conducted to investigate the variability when assessing mitochondrial respiration and content in human skeletal muscle.

Experiment 1: To examine the variability of two chambers from the same Oxygraph-2k respirometer, yeast cell suspension was used to measure the oxygen consumption rate.

Experiment 2: Mitochondrial respiration data of resting human skeletal muscle samples (n=76) from previously published studies [6, 9] were re-analyzed to estimate the variation between two chambers from the same high resolution Oxygraph-2k respirometer. Only mitochondrial respiration measurements with data from both chambers were used.

Experiment 3: Mitochondrial respiration data of resting human skeletal muscle samples (n=51) from unpublished studies were re-analyzed to estimate the within-sample variation when measured in triplicate using three chambers of a high resolution Oxygraph-2k respirometer (*i.e.*, across two respirometers). For each sample, the maximal mitochondrial respiration (CI + CII_p_) was calculated in two ways: the mean of the values from all three chambers (“All Three Values”), and the mean of the values after excluding those outside four times the CV of yeast respiration from the mean of three values (“Values Within 4× CV”). However, no more than one value was removed from the three repeated measurements of the same muscle sample; if more than one value were defined as outliers, only the value furthest away from mean was removed.

Experiment 4, 8 and 10: Resting human skeletal muscle samples were collected from 24 participants [*mean ± SD*, age: 24 ± 4 y; height: 179 ± 6 cm; mass: 78 ± 11 kg; V̇O_2 peak_: 46 ± 7 mL·min^-1^·kg^-1^] on two separate occasions (*i.e.*, Day 1 and 6). An additional sample was collected from 8 of the 24 participants [age: 24 ± 3 y; height: 177 ± 8 cm; mass: 79 ± 13 kg; V̇O_2peak_: 44 ± 10 mL·min^-1^·kg^-1^], providing three separate occasions (*i.e.*, Day 1, 6 and 11). Mitochondrial respiration (Experiment 4) and CS activity data (Experiment 8) were assessed to determine the CV and TE_M_ of the samples. These data were then used to calculated mitochondrial content-specific respiration (Experiment 10).

Experiment 5: Resting human skeletal muscle samples (n=72) collected from 24 participants [mean ± SD, age: 24 ± 4 y; height: 179 ± 6 cm; mass: 78 ± 11 kg; V̇O_2peak_: 46 ± 7 mL·min^-1^·kg^-1^] were used to measure CS activity. Each sample was loaded into three neighbouring wells on the same plate, from which the mean of the three wells was calculated. Any well with a value >10% from the mean of three wells was excluded from the calculation. Only samples measured in triplicate (*i.e.*, no wells were excluded from the calculation, n=42) were used to in the final analysis, calculate the variation between three technical repeats.

Experiment 6: Human skeletal muscle samples (n=55) collected before, 3 and 24 hours after an exercise intervention from 5 participants [mean ± SD, age: 28 ± 4 y; height: 175 ± 8 cm; mass: 70 ± 7 kg; V̇O_2peak_: 46 ± 10 mL·min^-1^·kg^-1^] were used to test the variation of the CS activity assay when analysing the same samples twice in the same day, approximately four hours apart, with the same reagents prepared in the morning. CS activity was measured using the same muscle lysate and protocol (*i.e.*, microplate). The CV and TE_M_ between microplates was then determined.

Experiment 7: Resting human skeletal muscle samples (n=72) collected from 24 participants [mean ± SD, age: 24 ± 4 y; height: 179 ± 6 cm; mass: 78 ± 11 kg; V̇O_2peak_: 46 ± 7 mL·min^-1^·kg^-1^] were used to test the variation of the CS activity assay when analysing the same samples twice, but with longer storage time and freshly made reagents each time. CS activity was measured using the same muscle lysate and protocol (*i.e.*, microplate) on two separate occasions, four weeks apart, each time using freshly made reagents. The CV and TE_M_ between microplates was then determined.

Experiment 9: CS activity values derived from two different protocols (*i.e.*, microplate vs cuvette) were compared, using data from unpublished studies. Resting human skeletal muscle biopsies (n=40) were collected from 10 participants [mean ± SD, age: 22 ± 5 y; height: 178 ± 11 cm; mass: 79 ± 12 kg; V̇O_2peak_: 47 ± 8 mL·min^-1^·kg^-1^]. The variation in results between the two protocols was then analyzed.

### Mitochondrial respiration using yeast cell suspension

Mitochondrial respiration in yeast cell suspension was measured using a published Oroboros reference assay with modification [52]. Commercial baker’s yeast powder was dissolved in sodium phosphate buffer containing 50 mM Na-PB, and adjusted to pH 7.1 at 30°C by pipetting repeatedly at the final concentration of 2 mg·mL^-1^. Mitochondrial respiration was measured in duplicate from 50 μL of yeast cell stock in 2 mL sodium phosphate buffer with 140,000 units·mL-1 catalase at 37°C using the high resolution Oxygraph-2k respirometer (Oroboros Instrument, Innsbruck, Austria). Glucose (20 mM) was used as the substrate to measure the NADH-linked respiration, and Azide (40 mM) was used to measure the non-mitochondrial oxygen consumption.

### Muscle biopsies

Muscle biopsies were performed under sterile conditions by a qualified, experienced, medical doctor. Local anaesthetic (Lidocaine, 1%) was injected into biopsy site and once numb, an incision (∼ 0.5 to 1.0 cm) was made through the skin and muscle fascia. Muscle was sampled using a suction-modified Bergström needle. One portion of the muscle sample (10–20 mg) was immediately immersed in 2 mL of ice-cold biopsy preservation solution (BIOPS) for measurements of mitochondrial respiration. The remaining muscle was cleaned of excess blood, fat, and connective tissue, and immediately frozen in liquid nitrogen and stored at −80°C for subsequent analyses.

### Mitochondrial respiration using permeabilized muscle fibres

Muscle fibres were mechanically separated using two pairs of forceps with sharp tips, in ice-cold BIOPS containing 2.77 mM CaK_2_EGTA, 7.23 mM K_2_EGTA, 5.77 mM Na_2_ATP, 6.56 mM MgCl_2_, 20 mM taurine, 50 mM MES, 15 mM Na_2_ phosphocreatine, 20 mM imidazole, and 0.5 mM dithiothreitol, adjusted to pH 7.1. Tissue permeabilization was carried out by gentle agitation for 30 min at 4°C in BIOPS containing 50 μg·mL^-1^ of saponin and was followed by three washes in MiR05, a respiration medium containing 0.5 mM EGTA, 3 mM MgCl2, 60 mM potassium lactobionate, 20 mM taurine, 10 mM KH2PO4, 20 mM Hepes, 110 mM sucrose, and 1 g·L^-1^ bovine serum albumin essentially fatty acid–free, pH 7.1 (27). Mitochondrial respiration was measured in duplicate (experiment 2) or triplicate (experiments 3 and from ∼2 to 4 mg wet weight of muscle fibres in the mitochondrial respiration medium (MiR05) at 37°C using the high resolution Oxygraph-2k respirometer (Oroboros Instrument, Innsbruck, Austria). Oxygen concentration (mM) and flux (pmol·s^-1^·mg^-1^) were recorded using DatLab software. Two different substrate-uncoupler-inhibitor titration (SUIT) protocols were used [44]. Our previously described substrate-uncoupler-inhibitor titration (SUIT) protocol [6, 9] was used for Experiment 2, whereas a modified version of this method (initial addition of 0.2 mM octanyolcarnitine and no determination of uncoupled respiration through complex IV) [53] was used for Experiments 3 and 4.

### Citrate synthase activity using a microplate

Approximately 20 mg of frozen muscle tissue was homogenized using a TissueLyser II (Qiagen, Valencia, CA) in an ice-cold lysis buffer (1:20 w/v) containing 50 mM Tris HCl, 150 mM NaCl, 1 mM EDTA, 5 mM sodium pyrophosphate; 1 mM sodium orthovanadate; 1% NP40, and a freshly added protease/phosphatase inhibitor cocktail (Cell Signaling Technology (CST), Danvers, MA, USA), adjusted to pH 7.5 with HCl. Homogenates were rotated end-over-end for 60 min at 4°C. Protein concentration was determined in triplicate using a Bradford assay (Bio-Rad protein assay dye reagent concentrate, Bio-Rad Laboratories, Hercules, CA), against bovine serum albumin standards (BSA, A9647, Sigma-Aldrich). The coefficient of variation for the protein assay was lower than 5%. CS activity was determined in triplicate on a microplate by adding the following: 5 µL of muscle homogenate (concentration ranging from 2 to 4 mg·mL^-1^), 40 µL of 3 mM acetyl coenzyme A, and 25 µL of 1 mM 5,59-dithiobis(2-nitrobenzoic acid) in Tris buffer to 165 µL of 100 mM Tris buffer (pH 8.3) kept at 30°C. After addition of 15 µL of 10 mM oxaloacetic acid, the plate was immediately placed in a plate reader at 30°C (xMark-Microplate; Bio-Rad), and after 30 s of linear agitation, absorbance at 412 nm was recorded every 15 s for 3 min. CS activity was reported as mol·h^-1^·kg^-1^.

### Citrate synthase activity using a cuvette

Analysis of CS acitivty in cuvettes was performed at the Mitochondrial Research Laboratory at the Murdoch Children’s Research Institute (Melbourne, Australia). CS enzyme activity was determined spectrophotometrically in post-600g supernatants of whole-muscle homogenates as previously described [54].

### Statistical analyses

All data in text, figures, and tables are presented as mean ± 95% confidence interval (CI), with exact *P* values presented, unless *P* < 0.001. For both mitochondrial respiration and CS activity, the coefficient of variation (CV) was calculated as the ratio of the SD of the values to the mean value. The technical error of the measurement (TE_M_) was calculated as previously described [19, 55], and relative TE_M_ is also represented as a percentage (calculated by dividing the TE_M_ value by the mean value). In Experiment 3, 4, 8, and 10, TE_M_ was calculated between three repeated measurements. Two value were randomly chosen from three repeated measurements of each sample and TE_M_ was calculated. This process was repeated ten times and the final TE_M_ was defined as the mean of ten calculated TE_M_. A paired t-test was used to assess differences in mitochondrial respiration or CS activity between chambers or samples in Experiment 1, 2, 4, 6, 7, 8, and 9. Paired or unpaired t-test was also used to assess the effect of muscle mass on mitochondrial respiration technical variability, and validate the outer mitochondrial membrane integrity test. One-way analysis of variance (ANOVA) was used to assess differences in mitochondrial respiration or CS activity between chambers or samples in Experiment 3, 4, 5, and 8. ANOVA was used assess the effect of absolute mitochondrial respiration value on technical variability, and validate the outer mitochondrial membrane integrity test. Pearson’s correlation coefficient was used to assess the relationship between the mitochondrial respiration measurements and CS activity measurements in Experiment 1, 2, 3, and 9. Lin’s concordance correlation coefficient and Bland-Altman plot were used to calculate the agreement between two methods measuring CS activity in Experiment 8. The simulations of the TE_M_ of the maximal respiration values (CI + CII_p_) to estimate the sample size are based on our previous method using the R software [19]. Statistical analyses were conducted using the statistical software package GraphPad Prism (V8.0), except the Lin’s concordance correlation coefficient and simulation analysis which were performed using the R software.

## DECLARATIONS

### Ethics approval and consent to participate

The study protocols were approved by the Victoria University Human Research Ethics Committee. The ethics numbers for the studies are HREC15-294, HRE17-075, HRE18-214, HRETH 11/94, and HRETH 12/189.

### Consent for publication

Written informed consent was obtained from the individuals involved in this study.

### Availability of data and materials

The datasets used and/or analyzed during the current study are available from the corresponding author on reasonable request.

### Competing interests

The authors declare that they have no competing interests.

### Authors’ contributions

JK, and DJB designed the study. JK, MJL and DJB drafted the initial manuscript. JK, NJS, JB, CG, ZW, XY, and JL conducted the study. NJS, JB, CG, ZW, XY, JL, and AJG contributed to the study design and provided feedback on the manuscript. All authors have read and approved the manuscript.

## Acknowledgements

The authors would like to thank Dr. Andrew Garnham and Dr. Mitchell Anderson for performing the muscle biopsies. The authors would also like to thank Adrienne Laskowski and Tegan Stait from Professor David Thorburn’s research group at Murdoch Childrens Research Institute for performing the CS activity assay using cuvette in Experiment 8. The authors would also like to thank Dr. Samantha Cassar, Charmaine DiQuattro, and Dr. Sudinna Hewakapuge for providing regular maintenance of the instruments. Lastly, the authors would like to thank all the participants in various studies.

## Notes

### Competing Interest Statement

The authors have declared no competing interest.

