## Supplemental tables for "Methodological considerations when assessing mitochondrial respiration and biomarkers for mitochondrial content in human skeletal muscle"

### SUPPLEMENTARY DATA

Supplementary Table 1: Summary of the coefficient of variation (CV) and technical error of measurement ($TE_M$) of mitochondrial respiration

| | No. of Participant | CV (%) | $TE_M$ (absolute) | Relative $TE_M$ (%) |
|---|---|---|---|---|
| Experiment 1: Two chambers (yeast) | 8 | 6.1 | 0.1 $nmol \cdot s^{-1} \cdot mg^{-1}$ | 6.3 |
| Experiment 2: Two chambers | 76 | 12.7 | 12.8 $pmol \cdot s^{-1} \cdot mg^{-1}$ | 15.4 |
| Experiment 3: Three chambers (All Three Values) | 51 | 16.7 | 15.2 $pmol \cdot s^{-1} \cdot mg^{-1}$ | 18.1 |
| Experiment 3: Three chambers excluding outliers (Values Within 4× CV) | 51 | 11.0 | 9.8 $pmol \cdot s^{-1} \cdot mg^{-1}$ | 11.7 |
| Experiment 4: Two repeated biopsies (Day 1 and 6) | 24 | 9.7 | 12.2 $pmol \cdot s^{-1} \cdot mg^{-1}$ | 15.0 |
| Experiment 4: Three repeated biopsies (Day 1, 6 and 11) | 8 | 12.8 | 12.1 $pmol \cdot s^{-1} \cdot mg^{-1}$ | 17.0 |
| Experiment 4: Two repeated biopsies (Day 1 and 11) | 8 | 13.5 | 9.8 $pmol \cdot s^{-1} \cdot mg^{-1}$ | 14.4 |

Supplementary Table 2: Summary of mean, standard deviation (SD) and 95% confidence interval (CI) of repeated muscle biopsies measurements

| | N | Mean | SD | 95% CI of mean |
|---|---|---|---|---|
| ***Experiment 4: Mitochondrial Respiration of three repeated biopsies (pmol·s$^{-1}$·mg$^{-1}$)*** | | | | |
| Day 1 | 24 | 81.4 | 21.7 | 72.3 – 90.6 |
| Day 6 | 24 | 80.6 | 21.0 | 71.8 – 89.5 |
| Day 1 | 8 | 71.1 | 12.9 | 60.4 – 81.9 |
| Day 6 | 8 | 77.1 | 20.1 | 60.3 – 93.9 |
| Day 11 | 8 | 64.9 | 16.0 | 51.5 – 78.2 |
| ***Experiment 8: CS Activity of three repeated biopsies (mol·h$^{-1}$·kg$^{-1}$)*** | | | | |
| Day 1 | 24 | 2.7 | 0.7 | 2.4 – 3.0 |
| Day 6 | 24 | 2.5 | 0.4 | 2.4 – 2.7 |
| Day 1 | 8 | 2.4 | 0.6 | 1.8 – 2.9 |
| Day 6 | 8 | 2.4 | 0.4 | 2.0 – 2.7 |
| Day 11 | 8 | 2.3 | 0.3 | 2.0 – 2.6 |
| ***Experiment 10: Mitochondrial Respiration/ CS Activity of three repeated biopsies (mol·h$^{-1}$·CS$^{-1}$)*** | | | | |
| Day 1 | 24 | 30.2 | 6.2 | 27.6 – 32.8 |
| Day 6 | 24 | 32.0 | 6.8 | 29.1 – 34.9 |
| Day 1 | 8 | 31.4 | 7.0 | 25.6 – 37.3 |
| Day 6 | 8 | 33.0 | 7.4 | 26.8 – 39.2 |
| Day 11 | 8 | 28.1 | 6.0 | 23.1 – 33.2 |

Supplementary Table 3: Summary of the coefficient of variation (CV) and technical error of measurement ($TE_M$) of CS activity

| | No. of Participant | CV (%) | $TE_M$ (absolute) | Relative $TE_M$ (%) |
|---|---|---|---|---|
| Experiment 5: Three technical repeats on the same plate | 42 | 3.5 | 0.1 $mol \cdot h^{-1} \cdot kg^{-1}$ | 4.1 |
| Experiment 6: Two repeated measurements in the same day (same muscle lysate) | 55 | 10.2 | 0.4 $mol \cdot h^{-1} \cdot kg^{-1}$ | 12.8 |
| Experiment 7: Two repeated measurements four weeks apart (same muscle lysate) | 72 | 30.5 | 0.7 $mol \cdot h^{-1} \cdot kg^{-1}$ | 30.9 |
| Experiment 8: Two repeated biopsies (Day 1 and 6) | 24 | 10.1 | 0.3 $mol \cdot h^{-1} \cdot kg^{-1}$ | 12.3 |
| Experiment 8: Three repeated biopsies (Day 1, 6 and 11) | 8 | 12.6 | 0.4 $mol \cdot h^{-1} \cdot kg^{-1}$ | 15.2 |
| Experiment 8: Two repeated biopsies (Day 1 and 11) | 8 | 14.1 | 0.4 $mol \cdot h^{-1} \cdot kg^{-1}$ | 18.5 |
| Experiment 9: Two protocols (microplate vs cuvette) | 40 | 17.0 | 1.3 $mol \cdot h^{-1} \cdot kg^{-1}$ | 21.9 |

Supplementary Table 4: Summary of the coefficient of variation (CV) and technical error of measurement ($TE_M$) of mitochondrial-specific respiration

| | No. of Participant | CV (%) | $TE_M$ (absolute) | Relative $TE_M$ (%) |
|---|---|---|---|---|
| Experiment 10: Two repeated biopsies (Day 1 and 6) | 24 | 12.5 | 5.5 mol·h$^{-1}$·CS$^{-1}$ | 17.6 |
| Experiment 10: Three repeated biopsies (Day 1, 6 and 11) | 8 | 19.0 | 7.1 mol·h$^{-1}$·CS$^{-1}$ | 23.0 |
| Experiment 10: Two repeated biopsies (Day 1 and 11) | 8 | 21.1 | 7.5 mol·h$^{-1}$·CS$^{-1}$ | 25.1 |
